# Donor specific transcriptomic analysis of Alzheimer’s disease associated hypometabolism highlights a unique donor, microglia, and ribosomal proteins

**DOI:** 10.1101/2019.12.23.887364

**Authors:** Sejal Patel, Derek Howard, Alana Man, Deborah Schwartz, Joelle Jee, Daniel Felsky, Zdenka Pausova, Tomas Paus, Leon French

## Abstract

Alzheimer’s disease (AD) starts decades before clinical symptoms appear. Low glucose utilization in regions of the cerebral cortex marks early AD and is clinically useful. To identify these regions, we conducted a voxel-wise meta-analysis of positron emission tomography studies that compared AD patients with healthy controls. This meta-analysis included 27 studies that assayed glucose utilization in 915 AD patients and 715 healthy controls. The resulting map marks hypometabolism in the posterior cingulate, middle frontal, angular gyrus, middle and inferior temporal regions. Using the Allen Human Brain Atlas, we identified genes with expression patterns associated with this hypometabolism pattern in the cerebral cortex. Of the six brains in the Atlas, one demonstrated a strong spatial association with the hypometabolism pattern. Previous neuropathological assessment of this brain from a 39-year-old male noted a neurofibrillary tangle in the entorhinal cortex. Using the transcriptomic data, we estimate lower proportions of neurons and more microglia in the hypometabolic regions when compared with the other five brains. Within this single brain, signal recognition particle (SRP)-dependent cotranslational protein targeting genes, which primarily encode cytosolic ribosome proteins, are highly expressed in the hypometabolic regions. Analyses of human and mouse data show that expression of these genes progressively increases across AD-associated states of microglial activation. In addition, genes involved in cell killing, chronic inflammation, ubiquitination, tRNA aminoacylation, and vacuole sorting are associated with the hypometabolism map. These genes suggest disruption of the protein life cycle and neuroimmune activation. Taken together, our molecular characterization of cortical hypometabolism reveals a molecular link to AD associated hypometabolism that may be relevant to preclinical stages.

## Introduction

Alzheimer’s disease, one of the most prevalent neurodegenerative diseases, is thought to affect approximately 5% of those aged 60 years and above worldwide [1]. It is the most common form of dementia, which is clinically characterized by a severe decline in cognitive functioning and defined neuropathologically by the emergence and topographical progression of amyloid plaques, neurofibrillary tangles, and neuronal loss[2].

Currently, fluorodeoxyglucose positron emission tomography (FDG-PET) is a primary frontline tool for the diagnosis of dementia and its subtypes. FDG-PET uses a radioactive tracer - [^18^F] fluorodeoxyglucose - to measure glucose metabolism within the brain [3], with altered cerebral glucose metabolism indicating AD with high sensitivity and specificity [4]. Importantly, patterns of hypometabolism can be seen in at-risk individuals decades before the development of symptoms [5–9]. This timing supports the concept that AD exists on a spectrum or continuum of pathologies that includes stages of subtle cognitive decline, mild cognitive impairment, and dementia [10,11]. Despite the clear link between metabolic changes measured by FDG-PET and risk for AD, it remains unclear which etiopathological mechanisms are responsible for driving these changes.

Using the Allen Human Brain Atlas, we sought to characterize the pattern of regional hypometabolism found in patients with AD. By integrating this atlas with a meta-analytic map of FDG-PET differences, we identified genes with spatial patterns similar to that of the lower glucose metabolism in the human brain. These genes that were associated with hypometabolism in the cerebral cortex were further tested for expression changes across AD-associated states.

## Materials and methods

### Meta analysis of Alzheimer’s FDG-PET studies

We performed a meta-analysis of FDG-PET studies that compared, at rest, Alzheimer’s patients with healthy controls. To compile a list of studies, a literature search was conducted on studies from January 1985 to January 2012. We used the following search query: [FDG-PET OR positron emission tomography OR fluorodeoxyglucose OR glucose metabolism] AND [dementia]. Studies were examined to fulfill the following criteria: (1) original research papers available in English (no case studies or reviews); (2) participants examined using [18F] FDG-PET at rest (no functional tasks); (3) AD patients compared with age-matched healthy controls; (4) clinical diagnosis of AD using NINCDS-ADRDA [12] or DSM-III [13] criteria; and (5) whole-brain analyses (no region-of-interest analyses) conducted in standardized stereotaxic space with available coordinates. Each article was read twice to determine if the study met the inclusion criteria.

Coordinates of regional hypometabolism peaks from retained articles were used to create ALE maps using BrainMap’s GingerALE application (www.brainmap.org/ale) [14]. This software assigns each voxel an activation likelihood estimate that is equal to the probability of at least one of the reported peaks of hypometabolism being located in that voxel [15]. To find distinct anatomical clusters, these voxelwise maps were clustered (min cluster extent = 500mm3; false discovery rate q= 0.05). The identified clusters were then used to determine a threshold that marks which samples are inside regions of hypometabolism.

### Gene expression data

The Allen Human Brain Atlas provides a comprehensive transcriptional landscape of the adult human brain [16]. The Atlas was obtained from six individuals (five males, one female), with age ranging from 24 to 57 years. Custom 64K Agilent microarrays were used to assay genome-wide expression in 3,702 spatially-resolved samples (232 named brain regions). We also used the RNA-sequencing datasets that were generated on the Illumina HiSeq2000 platform. These RNA-sequencing data were quantified with transcripts per million (TPM) and assayed a subset of anatomic structures from two of the six brains. The Allen Institute normalized the data and adjusted for array-specific biases, batch, and dissection method. Microarray probes were filtered using quality control data provided by Miller et al. [17]. After this filtering, 31,452 probes remained of the 58,692 on the microarray.

### Differential expression analyses

The microarray dataset was first used at the sample and donor level to identify genes that are differentially expressed in the regions of hypometabolism identified by the ALE-based analysis. Expression values were mean-averaged for genes with multiple probes, resulting in 15,143 genes. This analysis was restricted to samples from the cerebral cortex, as marked by the Allen Human Brain Atlas annotations (allocortical regions, namely the hippocampal formation and piriform cortex, were excluded). For each donor and gene, expression values were compared between samples inside and outside of the hypometabolic regions using the Mann-Whitney U test. The Allen Institute provided MNI coordinates, which were used to map expression values into the voxel space of the meta-analysis. For analyses that spanned multiple donors, Fisher’s method was used to generate a single meta p-value for each gene and direction [18]. We used the Benjamini–Hochberg false discovery rate (FDR) procedure for multiple test correction to adjust for the many tested genes [19].

### Gene Ontology enrichment analysis

The Gene Ontology (GO) provides gene-level annotations that span specific cellular components, biological processes, and molecular functions [20]. These annotations, defined by GO terms, were required to have annotations for 10 to 200 tested genes (6,333 GO groups annotating 14,241 unique genes). To test for enrichment, we sorted the genes from the most overexpressed to underexpressed in regions of hypometabolism. Within this ranking, the area under the receiver operating characteristic curve (AUC) was used to test for gene ontology terms that are enriched in either direction (overexpressed: AUC > 0.5, underexpressed: AUC < 0.5). The Mann-Whitney U test was used to determine statistical significance with FDR correction for the GO groups tested. We used GO annotations from the GO.db and org.Hs.eg.db packages in R, version 3.8.2, which were dated April 24, 2019 [21,22]. We used the REVIGO tool to summarize many terms that were significant after correction [23]. We used the default REVIGO parameters with uncorrected p-values for the input GO groups and restricted this analysis to the biological process branch of the GO.

### Estimation of Cell-Type Proportions

The Marker Gene Profile (MGP) tool was used to estimate cell-type proportions from the cerebral cortex expression profiles [24]. This method uses the first principal component of the expression of cell-type specific genes to estimate the relative abundance of a cell-type. We used 21 top marker genes obtained from a single cell study of the adult human temporal cortex [Supplementary Table S3 in [25]]. This study used transcriptomic profiles to cluster cells into astrocyte, neuron, oligodendrocyte, oligodendrocyte precursor, microglia and endothelial groups. These marker genes were used to calculate AUC values and estimate cell-type proportions with the MGP tool. Proportions were estimated separately for each donor across the same cortical samples used in the differential expression analysis.

### Single-cell RNA sequencing analysis of mouse microglia

Supplemental data from a single-cell RNA sequencing study of wild type and AD transgenic mouse model (5XFAD) were used to examine expression in immune cell types [26]. Keren-Shaul and co-authors profiled trancriptomically 8,016 immune cells from three wild type and three 5XFAD mice and clustered these cells into 10 distinct subpopulations based on expression. Of these 10 clusters, 3 expressed microglia markers. Two of these microglia clusters contained primarily cells from 5XFAD and not wild type mice and named them disease-associated microglia (DAM). For our analysis we consider these clusters separately as different microglial states: normal, intermediate (group II DAM), and AD associated (group III DAM).

### Single-nucleus RNA sequencing analysis

Supplemental data from a single-nucleus RNA sequencing study of the human prefrontal cortex were used to examine differential expression across AD states in microglia. Specifically, for each gene we extracted adjusted p-values (IndModel.adj.pvals), mean expression, and fold changes (IndModel.FC) from Supplement Table 2 in Mathys, Davila-Velderrain, et al. [27]. After quality control, Mathys, Davila-Velderrain, et al. clustered the transcriptomes of 70,634 nuclei from 48 individuals into eight broad cell-type clusters. For this work we focused on data from the 1,920 microglia nuclei. The 48 participants in this study were classified into no (24), early (15) and late (9) AD pathology. To test for enrichment of our genes of interest, we sorted the genes from the most overexpressed to underexpressed for the differential expression results for no versus early pathology and early versus late pathology analyses. Within this ranking, the area under the receiver operating characteristic curve measure (AUC) was used to test for genes that are significantly enriched in either direction. We also used the mean expression to determine which genes increase in expression across the three pathology groups. For a given set of genes, the hypergeometric test was used to determine if a greater number of genes increase across pathology than expected by chance.

## Results

### Meta-analysis of FDG-PET studies of AD

Our literature search for FDG-PET studies identified 3,229 titles. Screening of the abstracts yielded 230 relevant studies. Upon review of the full articles, 29 relevant studies remained. When two studies utilized the same patient population, one of the overlapping studies was excluded, resulting in a total of 27 studies yielding 33 independent samples with a total of 915 Alzheimer’s patients and 715 healthy controls (Supplement Table 1). Activation Likelihood Estimation (ALE) meta-analysis of these studies identified the following cortical regions as showing (consistently) lower glucose metabolism in patients vs. controls: posterior cingulate gyrus, middle frontal region, angular gyrus, middle and inferior temporal regions. A cluster analysis revealed 23 clusters (min cluster extent = 500mm3; FDR q= 0.05). A voxel-wise threshold of 0.006 was set to mirror this clustering map (Figure 1) and was used to determine if a given voxel was inside an AD-associated region of hypometabolism in subsequent transcriptomic analyses.

**Figure 1:**
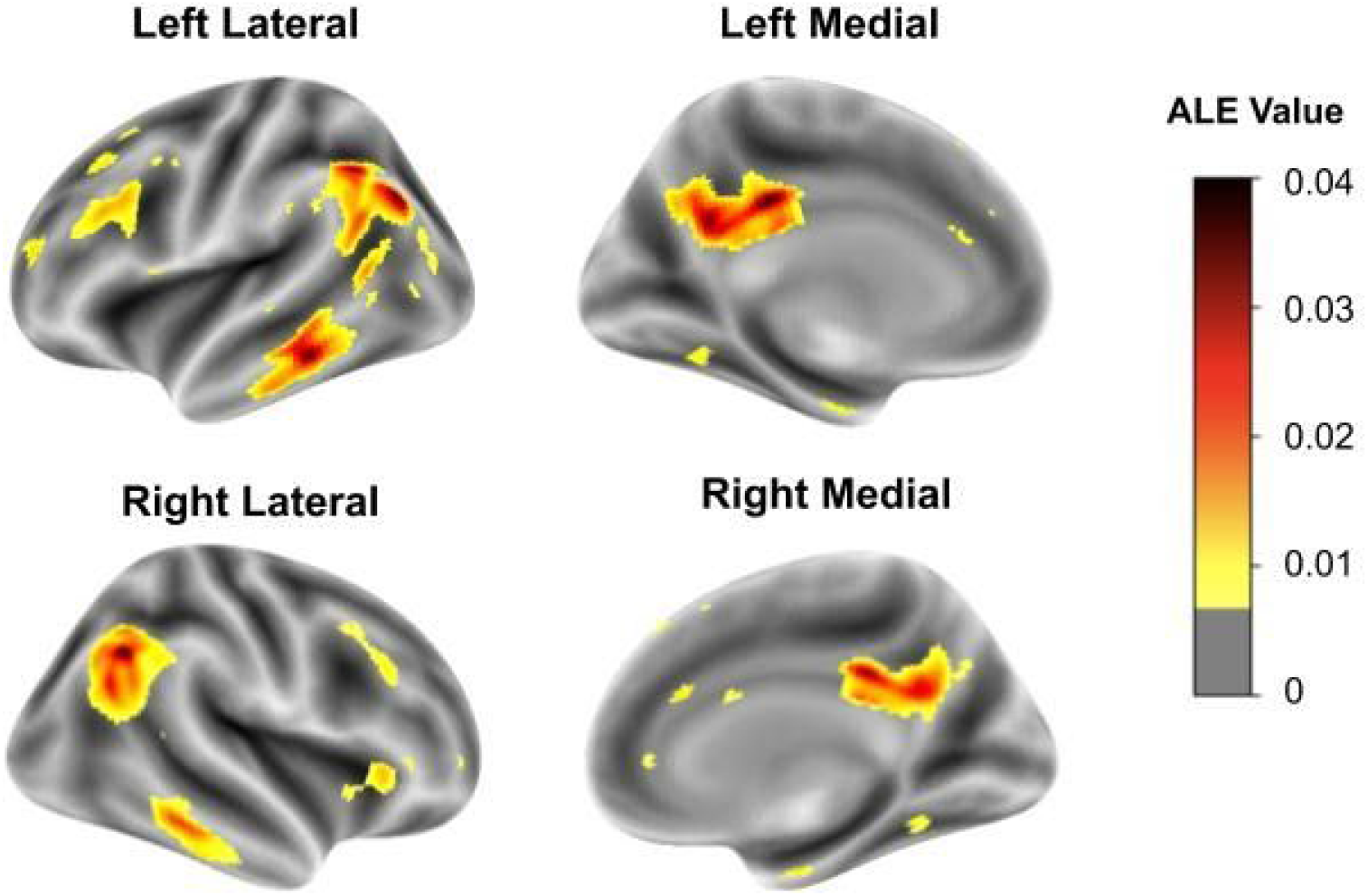
Cortical surface views of the ALE meta-analysis results. Regions where hypometabolism was not detected are transparent (ALE value of 0.006 or less). Lower glucose utilization (AD vs. controls) ranges from low (yellow) to high (black).

### Many genes are differentially expressed in cortical regions with AD associated hypometabolism

Using all six brains included in the Allen Atlas, we first identified the genes that were differentially expressed in the FDG-PET-defined hypometabolic regions of the cerebral cortex (1 female and 5 male, aged 24–57 years). The number of cerebral cortex samples that were profiled by the Allen Institute ranged from 182 to 481 per donor; of those, 5.9-9.9% overlapped with the hypometabolic regions. Of the 15,143 genes tested, 99 were significantly expressed at higher, and 51 at lower levels in these hypometabolic regions, after correction, across all donors. Substantial variability across the six brains in the Allen Human Brain Atlas has been previously noted both genome-wide and in the context of AD [28–31]. Given this variability, we then tested each brain separately. Strikingly, one brain drove the majority of the above atlas-wide signal for spatial expression overlap with the FDG-PET-derived map. In this brain (10021/H0351.2002), 647 genes were differentially expressed with 74% being expressed at lower levels in the hypometabolic regions (Supplement Table 2). In the remaining five donor brains, differentially expressed genes were only found in the oldest donor (donor 12876/H0351.1009, 57-year-old male). Taken together, our analysis of brain 10021/H0351.2002 marks it as an outlier with hundreds of genes that align spatially with the patterns of lower glucose metabolism observed in patients with AD (vs. controls).

### Brain specific analyses point to a unique donor

We examined the demographic information and metadata of this donor to help understand the above observation. Brain 10021/H0351.2002 was from a 39-year-old male African American individual. Postmortem interval was 10 hours, the lowest of the six donors. In agreement, RNA Integrity values (RIN) for this brain are higher than the other donors for all four of the regions assayed for RIN (frontal pole: 7.5, occipital pole: 7.1, cerebellum: 8.6, and brainstem: 7.3). As documented by the Allen Institute, this donor, like the others, had no known history of neuropsychiatric or neurological conditions. The presence of a broad range of drugs was tested for in postmortem blood by the Allen Institute. In donor 10021/H0351.2002, atropine, caffeine, lidocaine and monoethylglycinexylidide was detected at levels usually not toxicologically significant. We note that monoethylglycinexylidide is a metabolite of lidocaine, an anesthetic that is commonly used during dental procedures. Among the six donors, only 10021/H0351.2002 tested positive for lidocaine and monoethylglycinexylidide. The included brains were also classified as “normal” by a radiologist or pathologist. While considered neurotypical, it was noted that brain 10021/H0351.2002 contained a single neurofibrillary tangle in the entorhinal cortex. Neurofibrillary tangles in the hippocampus and entorhinal cortex are considered early events in AD progression [32]. Neurofibrillary tangles were not found in the other five brains (three of which are older than this donor). The presence of a neurofibrillary tangle is a unique feature of this individual and the postmortem interval and RIN values suggest tissue quality is not driving the Alzheimer’s associated molecular patterns that are observed.

### ER translocation genes are enriched for overexpression in areas of Alzheimer’s associated hypometabolism

To distil the molecular results we performed GO enrichment analysis on the transcriptome-wide results from donor brain 10021/H0351.2002. In total, 215 GO groups were significantly enriched (Table 1 shows the top 10 GO terms enriched for genes upregulated in hypometabolic regions). Due to the high degree of overlap in gene membership among our top GO terms, we used REVIGO tool to summarize them [23]. This tool removes redundant GO terms based on semantic similarity, providing a dispensability metric. Of the 98 biological process terms enriched for overexpression, three were assigned the lowest possible dispensability score of zero: SRP-dependent cotranslational protein targeting to membrane (GO:0006614, 87 genes, AUC = 0.874, p_FDR_ < 10^-28^), chronic inflammatory response (GO:0002544, 15 genes, AUC = 0.78, p_FDR_ < 0.05), and cell killing (GO:0001906, 94 genes, AUC = 0.60, p_FDR_ < 0.05). The strongest signal is from genes involved in SRP-dependent cotranslational protein targeting to membrane (Figure 2). This process targets protein translocation to the endoplasmic reticulum via the signal-recognition particle (SRP). These genes are primarily components of the cytosolic ribosome and henceforth referred to as ‘ER translocation’ genes. Six of these genes are found within the top 20 genes with higher expression in hypometabolic regions *(RPL34, RPL32, RPS27, RPS27A, RPL37A,* and *RPS15A*). In contrast, genes that are underexpressed in regions of hypometabolism are less significantly enriched for specific GO terms (lowest p_FDR_ = 7.3 × 10^-8^). However, these top terms contain more diverse themes (bottom half of Table 1), some of which have been previously implicated in AD. The most significant GO terms representing these themes are: ‘ubiquitin ligase complex’, ‘tRNA aminoacylation’, ‘ATPase activity, coupled’, ‘HOPS complex’ (involved in endosomal vesicle tethering), and ‘microtubule organizing center part’. The ubiquitin-proteasome system has been linked to AD [33]. Of the four genes that encode ubiquitin, three with available data are strongly overexpressed in regions of hypometabolism in this brain. In summary, this enrichment analysis points to spatial differences in vesicle fusion, protein translation, targeting, and degradation.

**Figure 2:**
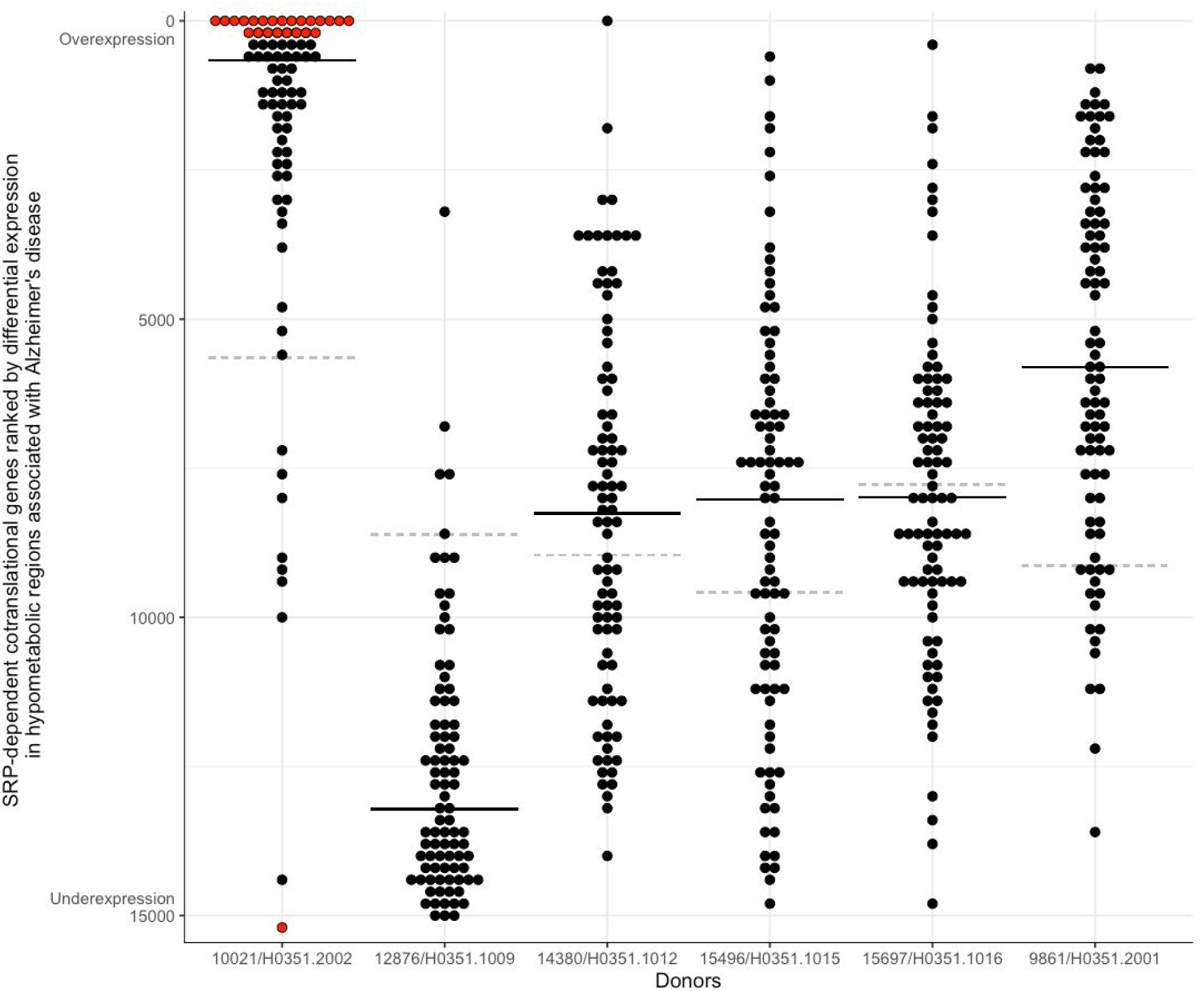
SRP-dependent cotranslational genes ranked based on differential expression in hypometabolic regions associated with AD. Genes are marked with dots, with the y-axis representing the genome-wide differential expression rank and ranges from overexpression (top) to underexpression (bottom). The black line marks the median expression rank of the SRP-dependent cotranslational genes. The dashed grey line marks the gene with the most stable expression between inside and outside of the hypometabolic regions for each donor. Red highlights genes that pass correction for multiple testing.

**Table 1:** Top GO groups enriched for differential expression in areas of Alzheimer’s associated hypometabolism in brain 10021/H0351.2002.

| Name | Genes | ID | AUC | p-value<sub>FDR</sub> |
| --- | --- | --- | --- | --- |
| SRP-dependent cotranslational protein targeting to membrane | 87 | GO:0006614 | 0.874 | 1.35E-29 |
| cotranslational protein targeting to membrane | 90 | GO:0006613 | 0.865 | 2.07E-29 |
| protein targeting to ER | 92 | GO:0045047 | 0.847 | 2.86E-27 |
| cytosolic ribosome | 87 | GO:0022626 | 0.856 | 3.45E-27 |
| establishment of protein localization to endoplasmic reticulum | 96 | GO:0072599 | 0.828 | 1.66E-25 |
| structural constituent of ribosome | 107 | GO:0003735 | 0.794 | 1.05E-22 |
| ribosomal subunit | 158 | GO:0044391 | 0.737 | 1.01E-21 |
| nuclear-transcribed mRNA catabolic process, nonsense-mediated decay | 104 | GO:0000184 | 0.783 | 2.07E-20 |
| protein localization to endoplasmic reticulum | 109 | GO:0070972 | 0.765 | 9.44E-19 |
| cytosolic large ribosomal subunit | 47 | GO:0022625 | 0.894 | 6.73E-18 |
| ..... |  |  |  |  |
| microtubule organizing center part | 145 | GO:0044450 | 0.395 | 0.00244 |
| DNA-dependent ATPase activity | 59 | GO:0008094 | 0.33 | 0.00145 |
| HOPS complex | 13 | GO:0030897 | 0.137 | 0.00135 |
| ATPase activity, coupled | 186 | GO:0042623 | 0.396 | 0.00026 |
| tRNA aminoacylation for protein translation | 40 | GO:0006418 | 0.268 | 9.84E-05 |
| amino acid activation | 43 | GO:0043038 | 0.275 | 8.24E-05 |
| aminoacyl-tRNA ligase activity | 33 | GO:0004812 | 0.243 | 8.24E-05 |
| cullin-RING ubiquitin ligase complex | 111 | GO:0031461 | 0.355 | 3.84E-05 |
| tRNA aminoacylation | 42 | GO:0043039 | 0.259 | 1.91E-05 |
| ubiquitin ligase complex | 195 | GO:0000151 | 0.368 | 7.35E-08 |

### Estimates of cell-type proportions are disrupted in hypometabolic regions in brain 10021/H0351.2002

To test if regional transcriptomic differences might be due to cell type proportions, we performed enrichment analyses of cell type-specific marker genes based on the differential expression results. In the five brains, microglia marker genes were expressed at low levels in the hypometabolic regions (underexpressed; AUC = 0.1, p_FDR_ < 10^-8^) while astrocyte and neuron markers were expressed at high levels (overexpressed; AUC > 0.66, p_FDR_ < 0.05). In contrast, brain 10021/H0351.2002 showed an opposite pattern of enrichment (Supplement Table 3). Using the Marker Gene Profile [24] tool, which uses a more complex parametric method, we also observe an interaction between hypometabolic regions and brain 10021/H0351.2002, whereby estimates of microglia proportions are higher inside hypometabolic regions in brain 10021/H0351.2002 (5 genes, t = 2.1, p = 0.033) and estimated proportions of neurons are lower (21 genes, t = −4.0, p < 0.0001).

### Validation of ER translocation genes with RNA sequencing data

Focusing on donor 10021/H0351.2002, the top-ranked gene ontology group, ‘SRP-dependent cotranslational protein targeting to membrane’/’ER translocation’, contains genes that are involved in the targeting of proteins to the endoplasmic reticulum. Given the high and ubiquitous expression of ribosomal genes, it is possible that the ER translocation signal is due to ceiling effects induced by the dynamic range of microarray gene expression profiling. To address this concern, we tested for the association using RNA sequencing data, which has a broader dynamic range. We again observe that the ER translocation genes are enriched (100 cerebral cortex samples, AUC = 0.733, p_FDR_ < 10^-9^). While limited in sample coverage for donor 10021/H0351.2002, the RNA sequencing data validates the finding of differential expression of ER translocation genes.

### ER translocation gene expression is high in AD-associated microglia (DAM)

Based on the differential expression of microglia markers in donor 10021/H0351.2002, we examined the ER translocation genes in microglia from an Alzheimer’s mouse model. We tested if the ER translocation genes increase in a stepwise pattern across the normal, intermediate, and full DAM clusters. For the 12,712 genes with data available, 6.5% monotonically increase in expression across these cell type clusters that represent distinct states. Of the 80 mouse homologs of the ER translocation genes, 75% increase in a stepwise fashion (Figure 3, hypergeometric p < 10^-52^). Compared with all gene ontology groups, this is the most significant enrichment (Supplement Table 6). In this single-cell dataset, ER translocation genes are expressed in AD associated microglia in a progressive pattern.

**Figure 3.**
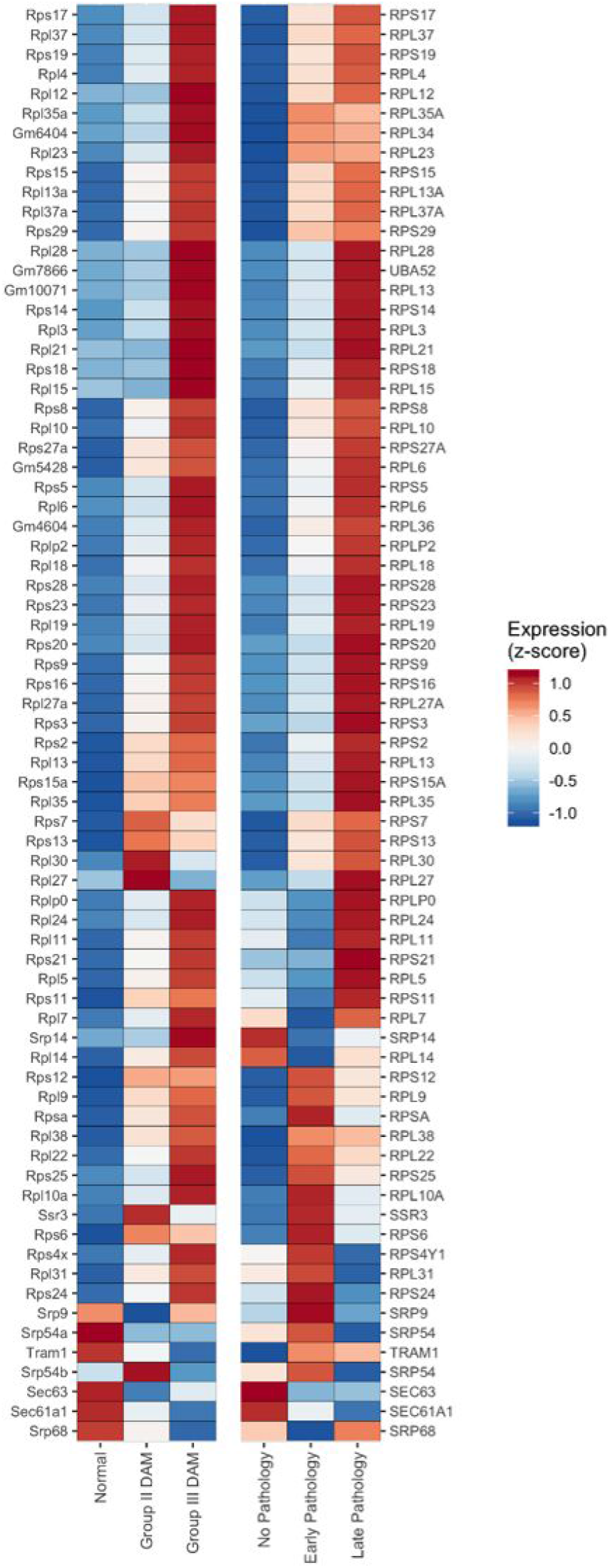
Heatmap of the ER translocation gene expression across three microglia cell clusters from AD mouse model (left half) and AD pathology subgroups (right half). Expression for each gene is z-scored with high expression in red and low in blue.

Genes are ordered based on hierarchical clustering using complete linkage (genes with similar expression across the mouse and human data are clustered together). Three human genes are duplicated because they have two homologous mouse genes *(RPL6, RPL13*, and *SRP54).* Human genes without homologous mouse genes are not displayed.

### Expression of ER translocation genes is correlated with AD pathology

Using data from a single-nucleus study of the human prefrontal cortex, we next tested if the ER translocation genes are differentially expressed across stages of AD pathology [27]. Guided by our findings in mice, we restricted our analyses to microglia. When comparing expression between no- and early-pathology subgroups we find that the ER translocation genes are enriched for higher expression in microglia from the early pathology individuals (Supplement Figure 1, 79 genes, AUC = 0.716, *p* <10^-10^). For the comparison between early- and late-pathology subgroups, the ER translocation genes are also enriched for higher expression in the late-pathology microglia (Supplement Figure 1, 77 genes, AUC = 0.627, *p* < 0.0005). Beyond these pairwise tests, we counted how many genes increase with disease progression. Broadly, for the 7,319 genes with data available, average microglial expression of 17.9% progressively increase across the pathological groups. For the ER translocation genes, this proportion triples to 55.8% (Figure 5, 43 of 77 genes progressively increase, hypergeometric p < 10^-13^). Compared to all GO groups, this is the second most significant group with the mostly overlapping set of cytosolic ribosome genes ranked first (Supplement Table 5). In this single-nucleus dataset, microglial expression of ER translocation genes is correlated with AD progression.

## Discussion

In this study, we projected the transcriptome of the cerebral cortex onto the spatial pattern of glucose hypometabolism found in AD cases. Of the six normal brains tested, only one demonstrated a strong spatial association between gene expression and the hypometabolism pattern. ER translocation genes, which encode proteins of the cytosolic ribosome and are involved in targeting protein translation to the endoplasmic reticulum, best align with the hypometabolic pattern in this brain. Using the transcriptomic data for this individual, we estimate a lower proportion of neurons and more microglia in hypometabolic regions. In support, prior neuropathological examination of this individual found a neurofibrillary tangle. Beyond this single brain, the same genes have a staged expression pattern that increases across cellular and pathological AD associated states in microglia.

It is striking that the ER translocation GO group was the most significantly enriched set in our analysis of the 10021/H0351.2002 donor brain and AD associated microglia. It is known that cytosolic ribosome genes are strongly co-expressed [35]. While we did not perform co-expression analysis, a change across this gene set will be easily detected with pathway or gene ontology analyses due to their high co-expression. This coherence is partly why it ranks above all other gene sets tested. Nonetheless, we note that a *RPL34* is a top ranked gene, providing a strong signal at the level of single genes. To gauge the chance of this GO group being top-ranked in multiple studies, we checked if the group is multifunctional or contains genes that are commonly differentially expressed. We found that this group ranked average in terms of multifunctional genes, relative to other groups (ranked 6,848th of 11,404 GO groups) [36]. In addition, this group was not top-ranked in any of the 635 studies systematically examined in a broad study of differential gene expression predictability [37]. More directly, the ER translocation genes are stable, with a below average prior probability of differential expression (ER translocation genes median = 0.246, remaining genes = 0.562, Mann-Whitney U test p < 10^-9^). Therefore, while ER translocation genes are strongly co-expressed, the prior likelihood of the ER translocation genes being differentially expressed is low.

It is plausible that brain atlases seeking to assay the normal brain may contain samples from donors who may be in the hypothetical stage of preclinical AD [34]. Our findings suggest that donor 10021/H0351.2002 may have been on this path. The ribosome and protein synthesis have been previously associated with mild cognitive impairment and AD (36–39). Pathological tau has also been shown to determine translational selectivity and co-localize with ribosomes (40, 41). Beyond the ER translocation genes, we note other GO groups with functional relevance. For example, ‘chronic inflammatory response’ and ‘cell killing’ genes were enriched for overexpression in the hypometabolic regions in brain 10021/H0351.2002. In the other direction, the genes in the homotypic fusion and protein sorting (HOPS) complex are underexpressed in hypometabolic regions in brain 10021/H0351.2002. This complex contains vacuole sorting genes and regulates autophagosome-lysosome fusion [38]. The top two most underexpressed gene sets in the hypometabolic regions are ‘ubiquitin ligase complex’ and ‘tRNA aminoacylation’. While ubiquitin ligase complex genes are underexpressed, genes encoding ubiquitin are overexpressed in the hypometabolic regions in brain 10021/H0351.2002. In summary, analysis of this single brain identifies genes that function in the protein life-cycle and neuroinflammation, which are known to be disrupted in AD [39–41].

## Declarations

## Acknowledgements

We thank Taylor Schmitz and Spiro Pantazatos for analyzing MRI scans. We thank the Allen Institute for Brain Science for creating the transcriptomic atlas of the human brain. We thank Ed Lein, Michael Hawrylycz, Jeremy Miller, Taylor Schmitz, and Shreejoy Tripathy for their insightful comments and suggestions.

## Authors’ contributions

DS, ZP, and TP designed and performed the FDG-PET meta-analysis. AM, JJ, SP, DH and LF performed the transcriptomic analyses. AM, SP, DH, DF, ZP, TP, and LF contributed to study design and the interpretation of the data. AM, SP, DH, DF, TP, and LF contributed to manuscript preparation. All authors read and approved the final manuscript and are accountable for their contributions.

## Funding

This study was supported by the CAMH Foundation, CAMH Discovery Fund, and a National Science and Engineering Research Council of Canada (NSERC) Discovery Grant to LF.

## Competing interests

LF owns shares in Cortexyme Inc., a company that is developing a gingipain inhibitor to treat AD. The other authors declare no conflict of interest.

## Ethics approval and consent to participate

Not applicable (all human data used are anonymized and in the public domain).

## Consent for publication

Not applicable (all human data used are anonymized and in the public domain).

## Availability of data and materials

Scripts, supplementary tables, and data files for reproducing the analyses are available online at https://figshare.com/articles/Donor_specific_transcriptomic_analysis_of_Alzheimer_s_disease_associated_hypometabolism_highlights_a_unique_donor_microglia_and_ribosomal_proteins/12_233552 and https://github.com/leonfrench/AD-Allen-FDG.

## Notes

https://figshare.com/articles/Donor_specific_transcriptomic_analysis_of_Alzheimer_s_disease_associated_hypometabolism_highlights_a_unique_donor_microglia_and_ribosomal_proteins/12233552

https://github.com/leonfrench/AD-Allen-FDG

### Summary of Updates

Simplified by removing associations with cholinergic and gingipain hypotheses

